# Pfs230 Domain 12 is a potent malaria transmission-blocking vaccine candidate

**DOI:** 10.1101/2024.11.09.622785

**Authors:** Maartje R. Inklaar, Roos M. de Jong, Dari F. Da, Lisanne L. Hubregtse, Maartje Meijer, Karina Teelen, Marga van de Vegte-Bolmer, Geert-Jan van Gemert, Rianne Stoter, Hikaru Nagaoka, Takafumi Tsuboi, Eizo Takashima, Cornelia G. Spruijt, Michiel Vermeulen, Roch K. Dabire, Emmanuel Arinaitwe, Anna Cohuet, Teun Bousema, Matthijs M. Jore

## Abstract

Malaria transmission-blocking vaccines (TBV) target sexual stage parasites that are transmitted to mosquitoes and critical for spread of the pathogen. The clinically most advanced TBV candidate contains part of the Pro-domain (Pro) and Domain 1 (D1) of *Plasmodium falciparum* surface protein Pfs230. Subunit vaccines that contain other domains of Pfs230 have so far failed to induce functional antibodies. Here, we produced eight single domain fragments of Pfs230 in *Drosophila melanogaster* S2 cells and assessed their immunogenicity in mouse immunizations. In addition to D1-specific antibodies, antibodies raised against Domain 12 (D12) showed strong recognition of Pfs230 on the surface of female parasites. Importantly, D12-specific antibodies showed strong functional transmission-reducing activity in membrane feeding assays with cultured parasites, an activity that was complement-dependent. Murine D12-specific antibodies further reduced mosquito transmission of parasites acquired from naturally infected parasite carriers. The D12 antigen was recognized by sera from an all-age cohort of individuals who had been naturally exposed to *Plasmodium falciparum* with antibody levels increasing with age. In conclusion, we identified Pfs230D12 as a promising new TBV candidate.

**Impact statement:** Pfs230 Domain 12 induces antibodies in mice that strongly reduce transmission of lab-cultured and naturally circulating malaria parasites to mosquitoes.

## Introduction

The burden of malaria has increased in recent years, with 450,000 fatal malaria cases in 2016 rising to 608,000 in 2022 ^1^. Malaria is caused by *Plasmodium* parasites, of which *Plasmodium falciparum* is the deadliest, that are transmitted by *Anopheles* mosquitoes. Transmission to mosquitoes relies on the uptake of sexual stage parasites, female and male gametocytes, through a bloodmeal. Inside the mosquito midgut, gametocytes activate to become female macrogametes and exflagellating male microgametes that fertilize to form zygotes. From this point onwards *Plasmodium* parasites continue their lifecycle by developing into ookinetes that traverse the epithelial layer and forming oocysts underneath the mosquito midgut epithelium. Within these oocysts, sporozoites are formed that find their way to the salivary glands, resulting in infectious mosquitoes. Transmission to mosquitoes forms a bottleneck in the lifecycle of malaria parasites and is therefore an attractive target for interventions. Transmission-blocking vaccines (TBVs) target this bottleneck with the aim to reduce the number of mosquitoes that become infectious; TBVs thereby form valuable assets for malaria elimination strategies ^2^.

TBVs induce antibodies in the human host against surface antigens of gametes, zygotes and/or ookinetes. These antibodies are taken up via the bloodmeal together with gametocytes and human complement. Inside the midgut, where parasites egress from the red blood cells and become accessible to antibodies, the antibodies prevent further development of the parasite through neutralisation or activation of human complement that results in parasite lysis. The functional activity of antibodies is commonly quantified by measuring the reduction in oocyst numbers compared to a negative control, and expressed as the percentage transmission-reducing activity (TRA). The clinically most advanced TBV candidate is Pfs230, which is essential for fertilization and further development into oocysts ^3^. Pfs230 is an abundant gamete surface protein that consists of fourteen 6-Cys domains (Figure 1A) ^4,5^. The large size of Pfs230 hampers recombinant expression of full-length Pfs230 and vaccine development has focused on expression of fragments of the protein ^6^. Immunization studies found that the Pfs230 Pro-domain (Pro) and Domain 1 (D1) induced a functional response in rodents ^7,8,9-12^, which formed the basis for the development of Pfs230D1-EPA, the only TBV candidate to date that progressed to phase 2 clinical studies ^13,14^. For many years, it has been unclear whether domains outside Pro and D1 contain epitopes for functional antibodies. Recent studies showed that functional monoclonal antibodies (mAbs) induced by whole parasite immunization or natural exposure target Pfs230 epitopes outside ProD1 ^15-18^. However, recombinant fragments containing non-ProD1 fragments of Pfs230, have so far failed to induce a functional response *in vivo* ^7-9^ and ProD1 thus remains the only Pfs230-based vaccine candidate with demonstrated *in vivo* efficacy described to date.

**Figure 1.**
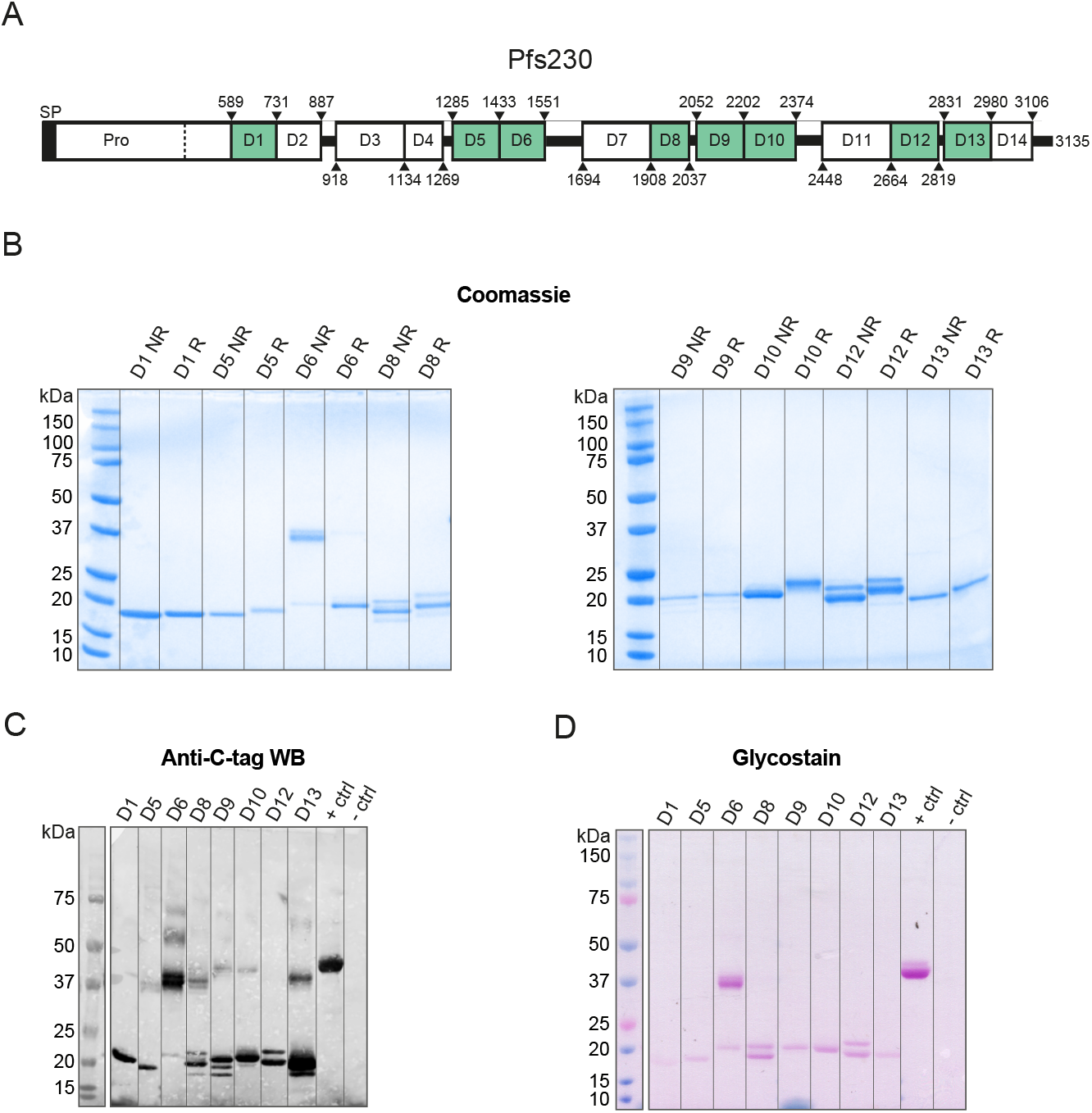
Recombinant single domain constructs produced in Drosophila melanogaster S2 cells. **(A)** Overview of full-length Pfs230. Predicted domain boundaries are indicated by amino acid numbers and are based on predictions made by Gerloff et al. ^5^. Single domain fragments with these boundaries were expressed in D. melanogaster S2 cells. Green colouring indicates successful expression of the fragment. (**B)** Coomassie-stained SDS PAGE gel of purified single domain fragments under reducing conditions (R) and non-reducing conditions (NR). **(C)** Western blot of recombinant fragments with α-C-tag antibody under non-reducing conditions. + and – controls are C-tagged protein and untransfected S2 cell line, respectively. **(D)** Glycosylation-stained SDS PAGE gel of recombinant fragments under non-reducing conditions. + and – controls are control proteins provided with the Pierce™ glycoprotein staining kit.

Here, we expressed all single Pfs230 domains in *Drosophila melanogaster* S2 cells that have been successfully used for expression of the 6-Cys domain protein Pfs48/45 ^19,20^. We obtained eight pure single domain fragments that were used to immunize mice. Of these fragments, Domain 12 (D12) induced antibodies with strong TRA in membrane feeding assays with lab cultured parasites and in membrane feeding assays with naturally circulating parasites from human donors. We also show that people with natural exposure to malaria parasites possess antibodies that recognize Pfs230D12. These results position Pfs230D12 as a promising new TBV candidate.

## Results

### Production of single domain Pfs230 protein fragments

We expressed single domain protein fragments with a C-terminal C-tag in *D. melanogaster* S2 cells (Supplementary tables 1-2). Nine of the fourteen constructs showed clear expression in S2 cell supernatants by western blot with an α-C-tag antibody (Supplementary figure 1A). The cell lines that showed clear expression, i.e. lines expressing D1, D3, D5, D6, D8, D9, D10, D12 and D13, were scaled up and proteins were purified using C-tag purification followed by size exclusion chromatography (Figure 1B). Unlike the other domains, D3 and D9 showed strong aggregation by size exclusion chromatography. Using the mild detergent Empigen® BB, we resolved aggregation for D9. D3 remained largely aggregated in the presence of Empigen® BB and was excluded from further analyses.

We obtained pure D1, D5, D6, D8, D9, D10, D12 and D13 proteins as determined by SDS-PAGE (Figure 1B) and western blot (Figure 1C) analyses. The proteins showed a slight change in apparent mass between reducing and non-reducing conditions, indicating that they form intramolecular disulphide bonds, as can be expected for 6-Cys domain proteins (Figure 1B, Supplementary table 2). D6 appeared as a dimeric protein on SDS-PAGE, which was resolved by the addition of reducing agent, indicating intermolecular disulphide bond formation (Figure 1B). All the antigens appear to be glycosylated, albeit to different extents (Figure 1D). Altogether, we obtained eight Pfs230 single domain antigens in sufficient quantity and purity for mouse immunizations.

### Pfs230D12 mouse antibodies recognize native Pfs230

For each selected Pfs230 protein construct, a group of five female mice was immunized three times with 20 μg antigen formulated in Montanide ISA720 and blood was collected 14 days after the third immunization (Figure 2A). A positive control group was immunized with 230CMB, a plant-produced protein containing the Pro-domain and D1, that was previously shown to induce strong TRA in rabbits ^21^. All mice generated antibody responses against the immunogen they were immunized with (Figure 2B, Supplementary figure 2). One mouse in the D12 group showed very low antibody responses (Supplementary figure 2) and sera from this mouse were therefore excluded from further analyses. Antibodies in pooled mouse sera recognized native Pfs230 in ELISA with gametocyte extract at different intensities (Figure 2C). Sera raised against D1, D5, D9, D10, D12 and D13 showed statistically significantly higher recognition compared to pre-immune sera. Recognition by sera against D6 and D8 was weaker and not statistically significant. We also tested recognition of native Pfs230 on the surface of purified live female gametes. Strikingly, only sera generated against 230CMB, D1 and D12 bound to the surface of female gametes (Figure 2D). Together, the results indicate that while sera raised against most single domain constructs recognize Pfs230 in parasite extract, only sera against D1 and D12 recognize Pfs230 on live female gametes.

**Figure 2.**
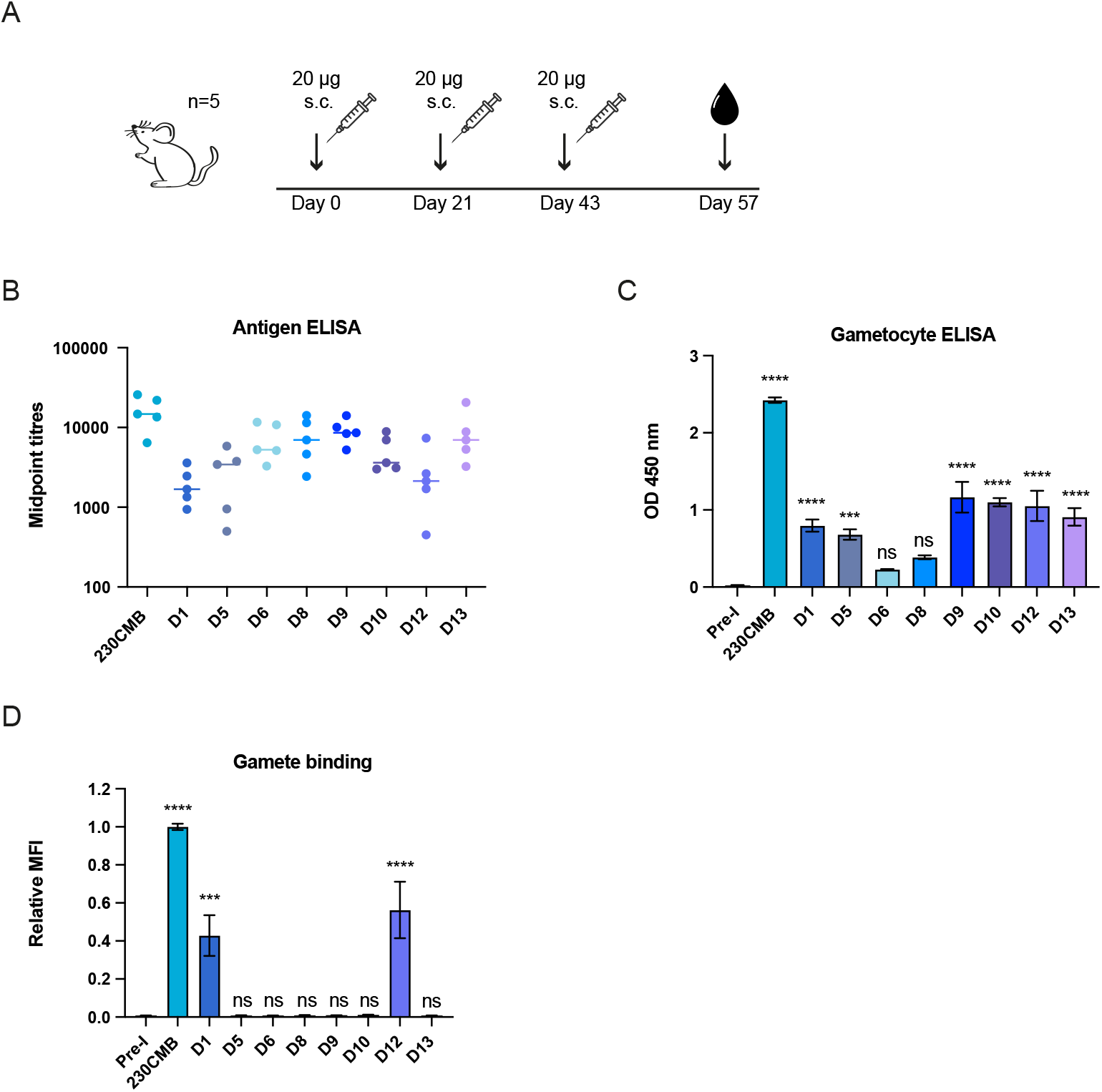
Antigen and parasite recognition by mouse antibodies raised against single Pfs230 domains. **(A)** Overview of mouse immunisation regimen. Groups of five mice were immunised subcutaneously (s.c.) with 20 μg antigen formulated in Montanide ISA-720. One group of mice was immunised with 230CMB (amino acids 444-730) ^21^ and was included as positive control. Final bleed sera were collected at day 57 for analyses of antibody responses in panels B-D. **(B)** Midpoint titers from antigen specific ELISA. Each dot represents an individual mouse and bars represent median values. **(C)** Gametocyte extract ELISA with pooled mouse sera, tested at 1:100 dilution. Values are means from two independent experiments with three technical replicates each and error bars represent s.e.m. Pre-I, pre-immune serum. **(D)** Female gamete binding assay with pooled mouse sera, tested at 1:40 dilution. Values are means (MFI= Mean Fluorescence Intensity) from two independent experiments with two technical replicates each, and error bars indicate s.e.m. The data was normalised against 230CMB in each independent experiment, to allow averaging across experiments. Statistical analysis in (C) and (D) done by comparing test groups to pre-immune group using ordinary one-way ANOVA with a Dunnett’s multiple comparison test using Pre-I as a reference (ns=not significant, ***=p<0.001, ****=p<0.0001).

### Pfs230D12 antibodies block transmission of Plasmodium falciparum NF54

To assess functional activity of the domain-specific antibodies, we performed standard membrane feeding assays (SMFA). In these assays, we allowed laboratory-reared *Anopheles stephensi* mosquitoes to feed on a mixture of cultured *P. falciparum* NF54 gametocytes and sera from immunized mice, and after 6-8 days we counted oocysts in the mosquito midgut to calculate TRA. In line with previous studies, sera raised against 230CMB and D1 showed strong TRA, reducing oocyst formation by 98.3% (95% CIs: 97.2-99.0) and 78.5% (95% CIs: 70.3-84.4) respectively (Figure 3A). Interestingly, D12 sera also showed strong TRA (95.2% TRA, 95%CIs: 93.1-96.6). Sera raised against other domains showed very low or no TRA, which is consistent with the binding assays where the antibodies in these sera failed to recognize the gamete surface (Figure 2D). Mass spectrometry confirmed purity of the D12 immunogen (Supplementary tables 3-4), and western blots with gametocyte extract and single domain fragments expressed with the wheat germ cell-free system further confirmed that the functional antibodies were D12-specific (Supplementary figure 3).

**Figure 3.**
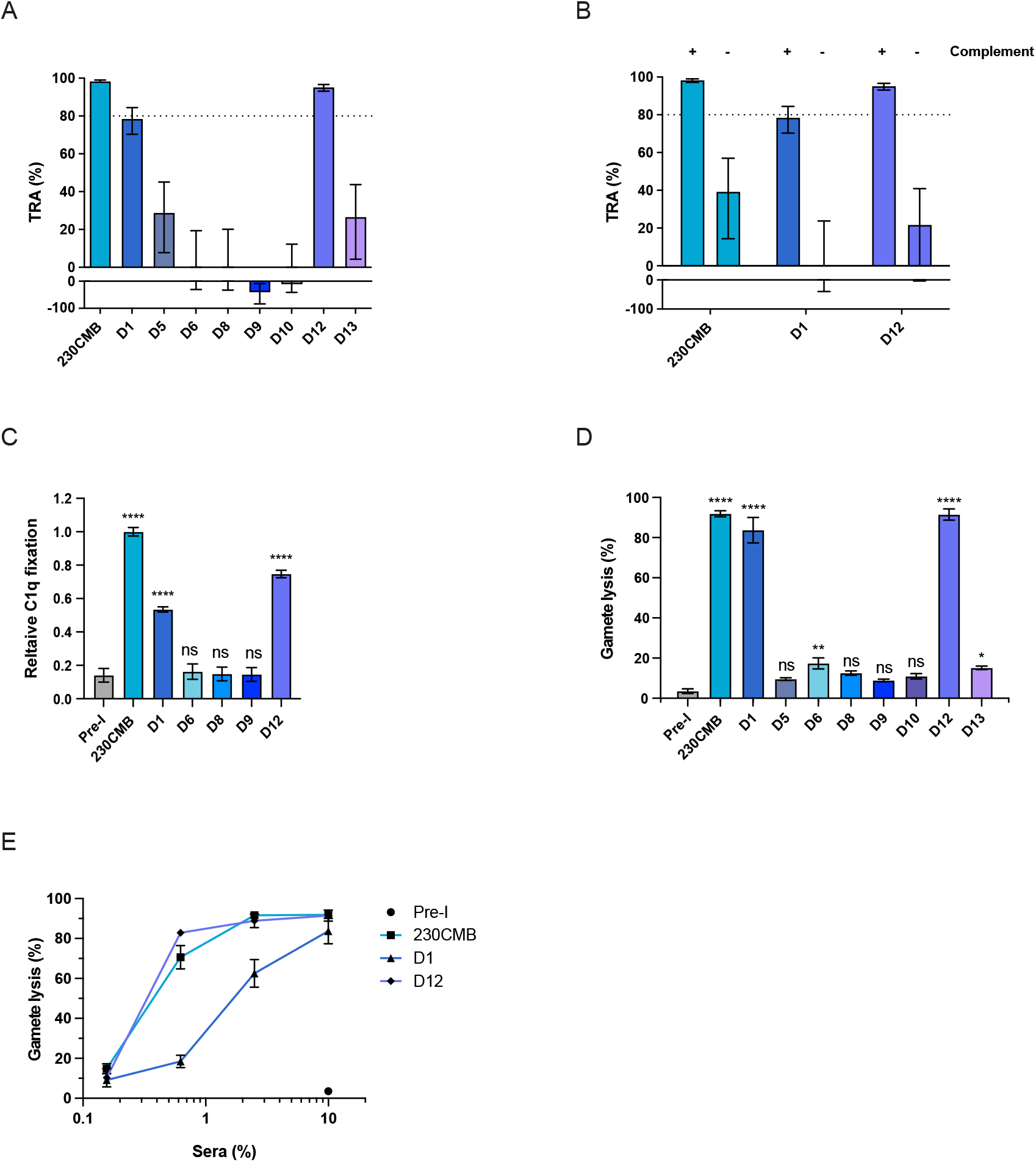
Functional activity of mouse antibodies raised against single Pfs230 domains. **(A)** Transmission reducing activity (TRA) of pooled mouse sera (day 57) in standard membrane feeding assay (SMFA) with cultured Plasmodium falciparum NF54 gametocytes and Anopheles stephensi mosquitoes. Values are estimates from two independent SMFA experiments with oocyst counts for 20 fully-fed mosquitoes per condition each, except for group D5 which had observations for 20 and 16 mosquitoes in two independent experiments, respectively. Dotted line indicates 80% TRA, which has previously been established as threshold for clinical development ^22^. Error bars indicate 95% confidence intervals. Pooled sera were tested at a final dilution of 1:9 in the presence of active human complement. (**B**) TRA of pooled mouse sera in SMFA, in the presence of active (+) or heat-inactivated (-) human complement. Pooled mouse sera were tested at a final dilution of 1:9. Values are estimates from two independent experiments with oocysts counts for 20 fully-fed mosquitoes per condition per experiment. **(C)** C1q deposition on the surface of female gametes in the presence of 2.5% pooled mouse serum, as assessed by flow cytometry. Values are means from two independent experiments with two technical replicates each and error bars indicate s.e.m. Data was normalised against 230CMB to allow averaging across experiments. Pre-I, pre-immune serum. (**D**) Female gamete lysis assay with 10% pooled mouse serum and active human complement. Values are means from two independent experiments with two technical replicates each and error bars indicate s.e.m. 0% lysis is defined as the number of live gametes present after incubation with human complement only. Statistical analysis in (C) and (D) was done by comparing test groups to pre-immune group using ordinary one-way ANOVA and accounted for multiple comparisons by Dunnett’s multiple comparison test (ns=not significant, *=p<0.05, **=p<0.01, ****=p<0.0001). **(E)** Pooled mouse sera that showed strong lysis in (D) were titrated in the gamete lysis assay. Values are means from two independent experiments with two technical replicates each and error bars indicate s.e.m.

Since the vast majority of functional Pfs230 antibodies described to date are complement-dependent, we tested sera against 230CMB, D1 and D12 in SMFA with either active or heat-inactivated human complement (Figure 3B). Sera showed substantial TRA only in the presence of active complement, demonstrating the complement-dependency of antibodies against these domains. To confirm that the complement-dependent activity is mediated by classical pathway activation, the ability of the mouse antibodies to fix C1q on the parasite surface was assessed in a flow cytometry assay with live female gametes. Antibodies against 230CMB, D1 and D12 were able to mediate C1q deposition, while antibodies against D6, D8 and D9 failed to do so (Figure 3C), in line with the SMFA results. We then tested whether the deposition of C1q leads to lysis of the female gamete in a flow cytometry lysis assay (Figure 3D-E). In this assay purified female gametes are incubated with mouse sera and human complement, and after incubation the percentage lysis is determined by live/dead staining. Sera against 230CMB, D1 and D12 showed over 80% lysis when tested at 1:10 dilution (Figure 3D). Other sera showed little, in most cases non-significant, lysis, which is consistent with results from the C1q deposition assay and SMFA. Titration of mice sera demonstrated similar potency for 230CMB and D12 sera, while D1 sera had slightly lower potency, similar to the trend observed in SMFA. Taken together, these data indicate that the D12 antigen can induce functional antibodies in mice capable of reducing transmission of lab-cultured malaria parasites to mosquitoes in a complement-dependent manner.

### Murine Pfs230D12 antibodies reduce transmission of naturally circulating gametocytes

To assess the functionality of D12 antibodies against naturally circulating gametocyte strains, we performed direct membrane feeding assays (DMFA) using blood of naturally infected gametocyte carriers from Burkina Faso. After removal of autologous plasma, red blood cells were mixed with pooled mouse sera and normal human serum containing complement, and fed to mosquitoes. After seven days oocysts were counted and TRA was calculated. We tested 230CMB and D12 sera that both showed strong TRA in SMFA, and further included pre-immune and D5 sera as negative controls. While D5 sera did not reduce oocyst formation, in line with SMFA results, the 230CMB and D12 sera reduced oocyst numbers across three independent experiments (Figure 4A-C). The estimated TRA values were 70.8% (95% CIs: 59.7-78.9) and 95.1% (95% CIs: 92.4-96.8) for 230CMB and D12 sera respectively (Figure 4D). The DMFA results thus show that D12 antibodies have strong TRA against naturally circulating gametocytes.

**Figure 4.**
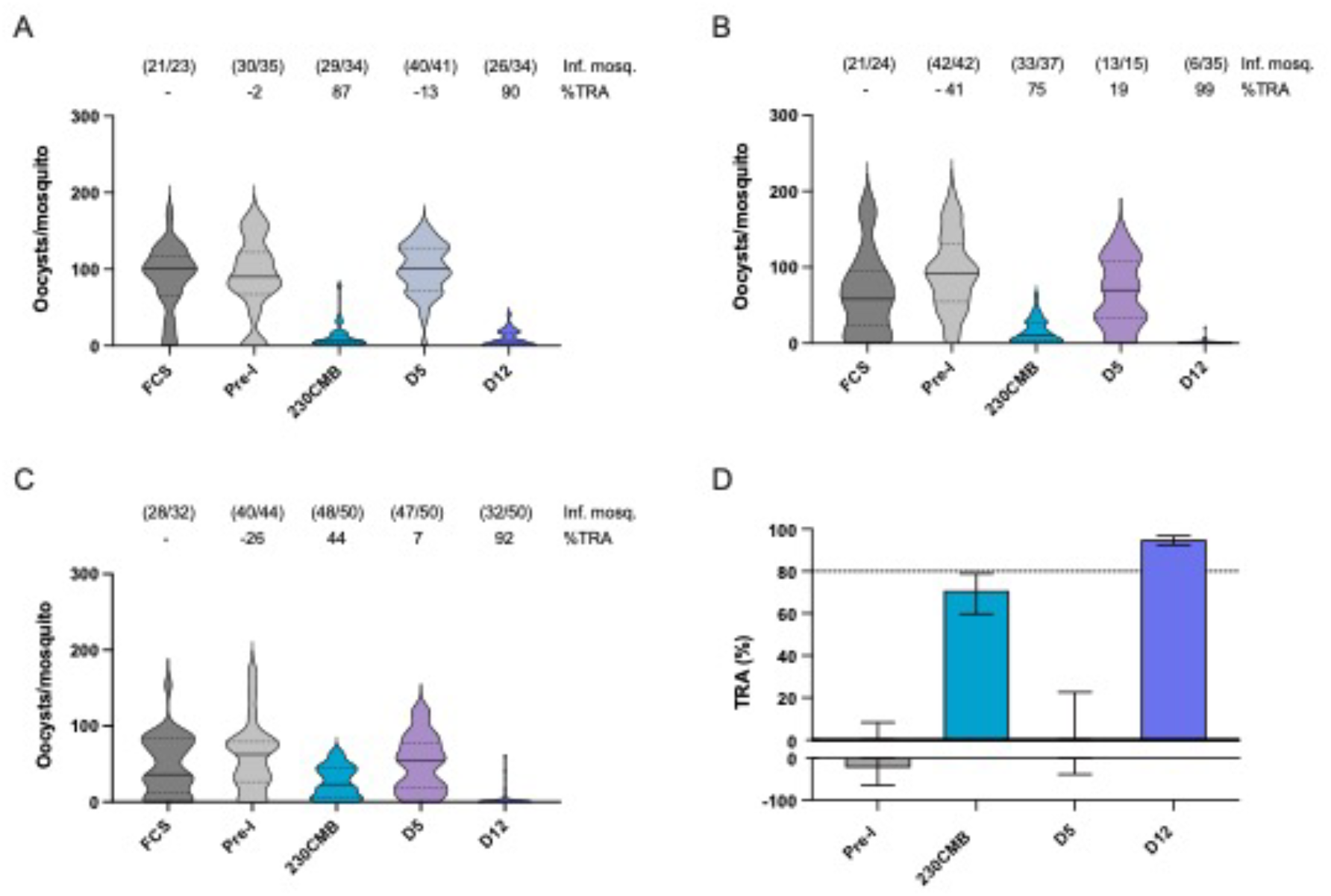
Transmission-reducing activity of Pfs230D12 sera in direct membrane feeding assay (DMFA) with naturally circulating gametocyte strains from volunteers in Burkina Faso. **(A-C)** Gametocytes from three volunteers were fed to Anopheles coluzzii mosquitoes, in the presence of mouse sera (day 57) and human complement. Pre-I, pre-immune serum. Data are shown as violin plots with the median indicated as a line and the interquartile range (IQR) (Q1 and Q3 quartiles) as dotted lines. Values above bars indicate percentage TRA, and the number of infected mosquitoes and the total number of mosquitoes between brackets (# infected mosquitoes/# total mosquitoes). **(D)** Estimate TRA values from three independent experiments (A-C) combined. Bars are estimated means and error bars indicate 95% confidence intervals. Dotted line indicates 80% TRA, which has previously been established as threshold for clinical development ^22^.

### Pfs230D12 is recognized by sera from individuals naturally exposed to P. falciparum

To assess natural antibody responses to D12, we screened plasma samples from a cohort of individuals residing in Tororo, an area in eastern Uganda where, at the time of sampling (2013-2017), transmission intensity was intense and perennial ^23^. Children ≤ 10 years of age and adults were eligible for enrolment. Purified IgG samples of these cohort participants were tested for transmission-reducing immune responses in the SMFA, as described above, as part of a larger study on naturally acquired transmission-reducing immunity. We detected significantly higher antibody levels in Ugandan samples compared to naïve control samples from Dutch donors (Mann-Whitney test, p < 0.0001) (Figure 5A). 139 out of 189 (73.2%) individuals were seropositive for D12. Antibody intensity was significantly higher in adults compared to school aged children (5-10 years old) and younger children (<5 years old) (Kruskal-Wallis test, p <0.0001) (Figure 5B). We observed a statistically significant correlation between antibody responses against 230CMB and D12 (Spearman’s, ρ = 0.4073, p < 0.0001) (Figure 5C). Whilst numbers were too small for meaningful statistical comparisons, we observed very strong D12 responses in two individuals whose total IgG isolated from plasma showed strong TRA in SMFA (Figure 5D). Together, we found that individuals living in a malaria-endemic country can generate antibodies against D12 and that antibody levels increase with age.

**Figure 5.**
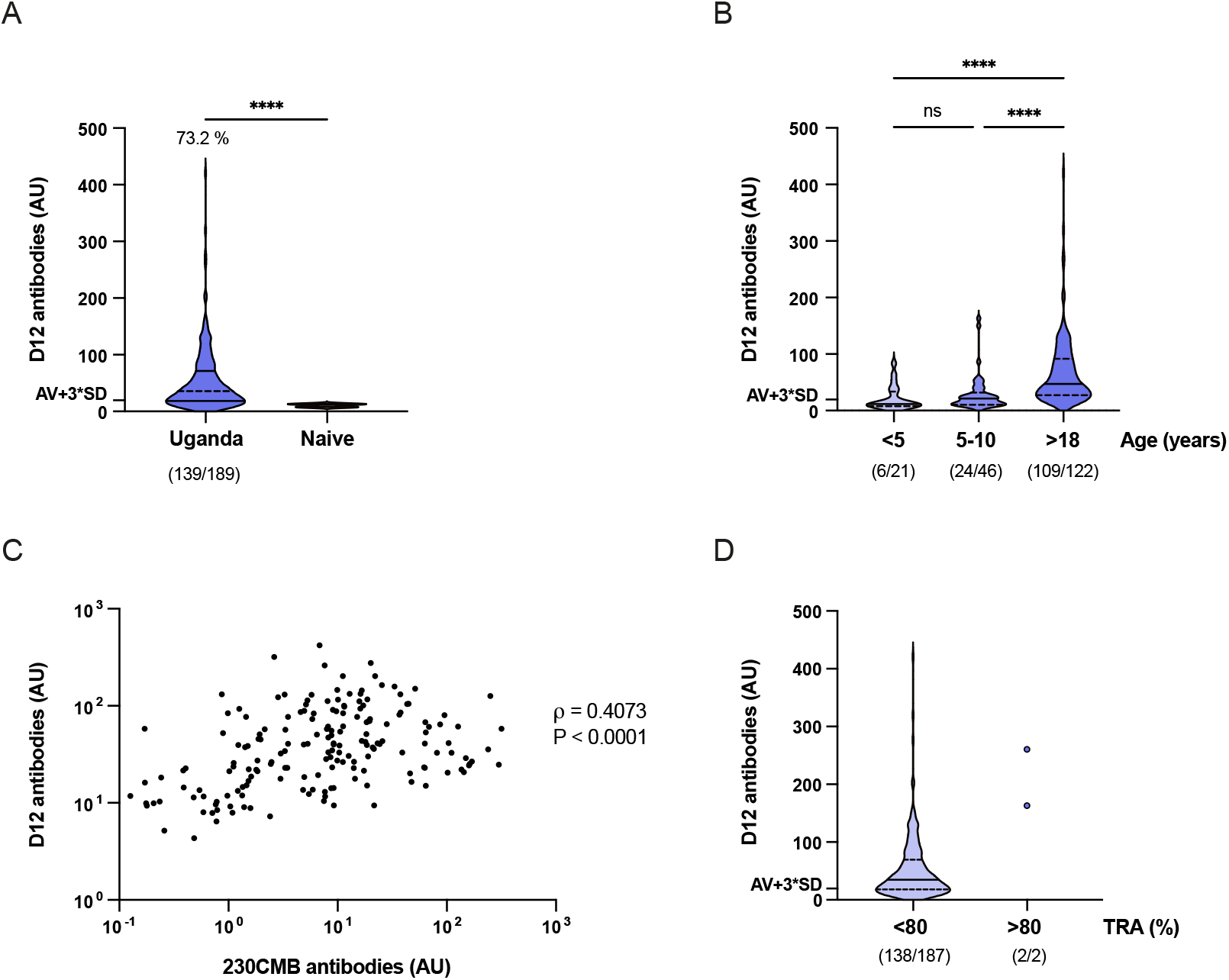
Recognition of Pfs230D12 by plasma from volunteers naturally exposed to Plasmodium falciparum. **(A)** Antibody levels against D12 in Ugandan plasma samples, as determined by ELISA. AU are arbitrary units calculated using a highly reactive plasma pool from Tanzania as reference. Naïve samples are pooled plasma samples from malaria-naïve Dutch donors (n=8). Groups were compared by one-sided Mann-Whitney test (****=p<0.0001). The threshold for positivity is marked and defined as the mean of naïve controls plus three standard deviations (AV + 3*SD). Percentage of positive samples is indicated above the graph. The number of positive samples and the total number of samples are depicted in between brackets (# positve samples/# total samples). **(B)** Antibody levels stratified by age. Groups were compared by Kruskal-Wallis test with Dunn’s correction for multiple testing (ns=not significant, ****=p<0.0001). Number of samples per age group are shown below the graph. **(C)** Correlation between antibody levels against D12 and 230CMB (Spearman’s ρ = 0.4073, p < 0.0001). Arbitrary units against 230CMB are calculated using serum from an individual that had high levels of antibodies against D12 (Donor A ^24^). Five individuals with 230CMB-specific antibody levels below 0.1 AU are not shown on graph. **(D)** Antibody levels stratified by TRA. Number of antibody positive over total tested individuals per group are shown below the graph. Data in (A), (B) and (D) are shown as violin plots with the median indicated as a line and the interquartile range (IQR) (Q1 and Q3 quartiles) as dotted lines.

## Discussion

Here, we expressed eight individual domains of Pfs230, a *P. falciparum* protein that is essential for parasite transmission and forms the basis of current malaria TBV development. Antibodies raised against the D12 domain showed potent binding to live female gametes *in vitro* and have strong functional TRA. Functional activity was demonstrated against lab-cultured gametocytes and gametocytes from naturally-infected parasite carriers from Burkina Faso. Furthermore, sera from Ugandan donors naturally exposed to *P. falciparum* showed immune recognition of D12 in an age-dependent manner. Taken together, our results identify Pfs230D12 as a promising new TBV candidate.

TBVs could be valuable tools for the elimination and eradication of malaria. Several TBV candidates have been identified, of which the ProD1 fragment of Pfs230 has progressed furthest in terms of clinical testing. This study aimed to comprehensively examine constructs outside ProD1 and is, to the best of our knowledge, the first to identify a non-ProD1 fragment of Pfs230 that can elicit functional antibodies. Two earlier immunization studies included fragments containing D12, produced in *Escherichia coli* ^8^ and wheat germ cell-free system ^9^. These constructs induced antibodies that recognized native Pfs230 on gametes, but the antibodies did not reduce transmission to mosquitoes _8,9_. This is in sharp contrast with our results that show high TRA for mouse antibodies raised against D12 produced in *D. melanogaster* S2 cells. It is likely that the expression system plays an important role with *D. melanogaster* S2 cells producing (more) properly folded D12 that is critical for raising functional antibodies (Figure 1B). This is in line with preclinical studies with the other 6-Cys family protein Pfs48/45 that showed that the host expression system and proper conformation of the antigen are essential for inducing functional responses ^25^.

While antibodies raised against the D12 antigen showed strong gamete recognition and functional TRA, we did not observe functional responses against D5, D6, D8, D9, D10 and D13 (Figure 3A). Strikingly, antibodies against most of these domains recognized native Pfs230 in gametocyte extract, but did not recognize Pfs230 on live gametes (Figure 2C-D) suggesting that epitopes on these domains are occluded by other parasite surface proteins that interact with, or are in close proximity of, Pfs230. Alternatively, epitopes on these domains may be close to the parasite membrane and therefore not accessible to antibodies. However, we cannot rule out that these domains of Pfs230 do contain functional epitopes, as the functional epitopes could have been absent in our recombinant antigens due to misfolding or masking by glycosylation in the *D. melanogaster* S2 cell expression system (Figure 1C).

Most if not all functional Pfs230 antibodies described to date are dependent on complement ^4,12,13,15,26-28^. The antibodies we raised against D12 are no exception to this rule. The complement-dependency of the D12 antibodies was shown in membrane feeding assays where antibodies lacked TRA in the presence of heat-inactivated complement (Figure 3B). The D12 antibodies can activate the classical pathway by fixing C1q (Figure 3C) and induce complement-mediated gamete lysis *in vitro* (Figure 3D and E), which is the presumed effector mechanism in the mosquito midgut.

The viability of D12 as vaccine candidate depends on several factors. In general, vaccine candidates should target conserved functional epitopes to generate cross-strain protection, should ideally be able to induce highly potent antibodies so lower overall antibody responses are needed for protection and should be immunogenic in humans. Like other sexual stage *Plasmodium* proteins, Pfs230 is well conserved; D12 contains a similar number of non-synonymous single nucleotide polymorphisms as the leading TBV candidate ProD1 ^16^. The D12-specific mouse antibodies not only block transmission of the reference strain *P. falciparum* NF54, but also block transmission of naturally circulating gametocytes that may be genetically diverse (Figure 4). However, whether the functional antibodies indeed target conserved epitopes on D12 is currently unknown and should be the focus of future research. This could be addressed by isolating and characterising monoclonal antibodies, either from immunized animals or from human donors with naturally acquired immunity, and assessing their structure-function relationships as recently done for ProD1 ^26,27,29^. Furthermore, we observed that many individuals in a Ugandan cohort were seropositive for D12, demonstrating that the D12 antigen is immunogenic in humans and that vaccine-induced antibody levels may be boosted by natural exposure and *vice versa*. Naturally acquired transmission reducing immunity is a rare phenomenon, at least at high levels of TRA that can be reproducibly demonstrated in the SMFA ^24^. Whilst now allowing for a formal assessment of a possible role of D12-specific antibodies in naturally acquired TRA, we made use of a larger cohort study where plasma samples were available alongside TRA estimates. Two individuals with high levels of TRA also had high levels of antibodies against D12. The strong correlation between anti-D12 antibodies and anti-Pfs230CMB antibodies makes it impossible to determine whether D12 antibodies were causally responsible for TRA. Purification of D12-specific antibodies from plasma and assessment of their TRA in SMFA ^24^ would allow us to demonstrate causality but this was not possible with the plasma volumes available. Nevertheless, our findings demonstrate that D12 antibodies are naturally acquired in an age-dependent manner – probably reflecting cumulative exposure to *P. falciparum* gametocytes – in a manner that is similar to antibodies to other Pfs230 domains and that individuals with naturally acquired TRA can have high levels of D12 antibodies.

In our immunisation studies we included 230CMB ^21^, containing ProD1, as positive control. It is encouraging to see that our non-optimized D12 immunogen induced similar levels of TRA and gamete lysis activity as ProD1. Future studies should assess whether the D12 antigen could be further optimized to induce stronger functional responses, for instance through coupling to carrier proteins, screening of glycosylation mutants, adjuvant screening and/or designing immunogen variants that are more stable or have improved epitope display. It will also be interesting to determine whether antibodies against D12 are synergistic or additive with antibodies against other TBV candidates such as Pfs230D1-EPA, as combining antigens may be an attractive approach to increase overall vaccine efficacy.

This study has several limitations. We aimed to assess all single domains of Pfs230 in our immunisation study, but could not produce six of the fourteen Pfs230 domains. These included D4 and D7, which have been shown to be targets for functional mAbs raised against whole parasites ^16,17^. Whether these domains can be produced recombinantly to induce functional antibodies thus remains unclear.

Furthermore, one of the mice in the D12 group showed very low antibody responses to the immunogen (Supplementary figure 2). Whether this low response is linked to the immunogen itself, or whether this was caused by external factors is currently unclear and should be assessed in future immunisation studies. Finally, we could test the mouse sera in only two DMFA experiments due to logistical challenges. The results from these DMFA experiments should therefore be interpreted with caution and warrant more in-depth studies to assess cross-strain protection.

In conclusion, our work shows how employing a different expression system to produce recombinant domains of Pfs230 can lead to the identification of a new malaria TBV candidate and that Pfs230D12 is a promising new candidate for further preclinical investigation.

## Materials and methods

### Protein construct design

Expression plasmids were created for all fourteen domains of Pfs230 (Figure 1A). Boundaries of the domains were based on previous research ^5^ and the linkers between domains were excluded. Sequences were codon optimized for expression in *Drosophila melanogaster* and synthesized (BaseClear). The Pfs230 single domains were cloned with an N-terminal BiP signal peptide, His_6_-tag and Alanine-serine linker, and a C-terminal glycine-serine linker followed by a C-tag into the pExpreS2.2 plasmid (ExpreS2ion Biotechnologies), downstream of the Actin+HSP70 promoter. The plasmids were verified by Sanger sequencing (Baseclear). The sequences of the inserts can be found in Supplementary table 1.

### Drosophila melanogaster S2 cell transfection and culture

The *D. melanogaster* S2 cell line (ExpreS2ion Biotechnologies) was used for the expression of all Pfs230 protein constructs. S2 cells were cultured in shake flasks with vented cap in EX-CELL420 media (Sigma-Aldrich), supplemented with 1% penicillin and streptomycin, at 25°C shaking 115 rpm. Cells were counted twice a week and resuspended to 8×10^6^ cells/mL alternately by dilution or centrifugation. For transfections, 2.5 mL cell suspension was mixed with 6.25 μg plasmid DNA and 25 μL ExpreS2 Insect-TR 5x transfection reagent (ExpreS2ion Biotechnologies) in a T12.5 T-flask. The transfected cells were then incubated at 25°C, and 1 mL of FBS was added after 3 hours. 4000 μg/mL geneticin was added as a selection agent after 24 h. Approximately 26 days after the transfection, the cultures were scaled up to shake flasks. During this step FBS and geneticin were removed by centrifugation and resuspending cells in EX-CELL420 to 8×10^6^ cells/mL. Supernatant was harvested 5 days after the cells were diluted, for protein expression analysis on western blot and protein purification.

### Protein purification

The S2 cell supernatant was concentrated from 200-300 mL to approximately 50 mL using the Masterflex EasyLoad (Masterflex). The Pfs230 single domain protein constructs were affinity purified with CaptureSelect C-tagXL pre-packed columns (Thermo Scientific) on an ÄKTA start (Cytiva), using 20 mM Tris wash buffer (pH 7.4) and 20 mM Tris + 2 M MgCl_2_ (pH 7.4) elution buffer. Peak fractions from the chromatogram were pooled and dialysed overnight in PBS. The sample was filtered and concentrated to approximately 600 μL. Subsequently, the sample was further purified using a Superdex75 10/300 GL column (Cytiva) with filtered and degassed PBS as running buffer. For Pfs230D9 and Pfs230D3 0.2% Empigen® BB (Sigma-Aldrich) was added to all purification buffers to decrease aggregation of the proteins. Superdex fractions containing pure monomer protein were pooled and the protein concentration measured using a Nanodrop spectrophotometer (Thermo Scientific). Samples were frozen in liquid nitrogen and stored at -70°C.

### SDS-PAGE analysis

For analysis of proteins from S2 expression, samples were mixed with 4x NuPAGE LDS sample buffer (Invitrogen), heated at 70°C for 10 min before loading on a 4-20% bis-tris polyacrylamide gel (GenScript). In the case of purified protein 1 μg protein was loaded per condition and the Precision Plus Dual Color protein marker (Bio-Rad) was used as size standard. The gels were stained for 30 minutes using Instant Blue Coomassie Protein Stain (Abcam). To reduce disulphide bonds, a final concentration of 10 mM dithiothreitol (DDT) was added in the preparation of the sample.

For protein analysis with parasite extract, *P. falciparum* NF54 gametocyte extract was prepared as described previously ^30^ and diluted to the equivalent of 500,000 gametocytes per well. A final concentration of 10 mM dithiothreitol (DTT) was added for reducing conditions. Gametocyte extract and 230CMB ^21^ samples were mixed with 4× NuPAGE™ LDS sample buffer and heated for 10 minutes at 70°C before loading on a 4–20% Bis-Tris gel (GenScript). 20 ng 230CMB was loaded per well and the Precision Plus Dual Color protein marker (Bio-Rad) was used as size standard.

For analysis of proteins expressed in wheat germ cell-free system, purified proteins (described previously ^9,17^) were mixed with SDS-sample buffer and TCEP-HCl (Pierce™), and denatured at 37°C for 30 minutes before loading on a 12.5% PAGE Tris gel (ATTO, Tokyo, Japan). 0.5 μg of each protein was loaded per well and Precision Plus Protein All blue standard (Bio-rad) was used as the size standard.

### Western blot analaysis

For western blot analysis of proteins expressed in S2 cells, a positive control with C-tag (Pro-CS3-6C, kindly gifted by Susheel Singh, 36.3kDa) and negative control (Pf3D7_1306500C no C-tag, transfected in S2 cells) was included. The bis-tris gels were blotted on a 0.45 μm nitrocellulose membrane using the TurboBlot system (Bio-Rad). The membranes were washed in between steps with PBS supplemented with 0.05% Tween20 (PBST), blocked overnight in 5% skimmed milk PBS (mPBS) at 4°C, and incubated for 1 hour at room temperature (RT) with the CaptureSelect Biotin Anti-C-tag conjugate (1/1000, Cat. No. 7103252100, Thermo Scientific) in 1% mPBST. Thereafter the membranes were incubated with 1/2500 IRDye Streptavidin 680LT (Cat. No. 926-68031, LI-COR) in 1% mPBST for 1 hour at RT. The blot was developed with Clarity Max Western ECL substrate (Bio-Rad) and imaged with the Odyssey CLX (LI-COR).

For western blots with gametocyte extract, gels were transferred to a 0.45 μm nitrocellulose membrane using the Trans-Blot Turbo transfer system (Bio-Rad). The blots were blocked with 5% skimmed milk in PBS before incubation with 1/5000 polyclonal serum from mice immunized with Pfs230D12. After washing, the strips were incubated with 1/3000 diluted polyclonal rabbit anti-Mouse IgG HRP (Cat. No. P0260, DAKO). Blots were developed with Clarity Max Western ECL substrate (Bio-Rad) and imaged on the ImageQuant™ LAS 4000 (GE Healthcare).

For western blot analysis of proteins expressed in wheat germ cell-free system, the SDS-PAGE gels were transferred to an Amersham Hybond P Low fluorescence 0.2 μm PVDF membrane (Cytiva) using the Trans-blot SD semidry transfer cell (Bio-Rad) (25 V, 126 mA/gel, 75 minutes). The blots were blocked with 5% skimmed milk in PBST before incubation with 1/1000 polyclonal serum from mice immunized with Pfs230D12 (diluted in PBST) for 1 hour at RT and over night at 4°C. After washing, the blots were incubated with 1/10,000 polyclonal Sheep anti-Mouse IgG HRP (Cat. No. NA931VS, Cytiva). Blots were developed with Immobilon Western chemiluminescent HRP substrate (Cat. no. WBKLS0500, Millipore) and imaged on the LAS-4000 (FUJIFILM, Tokyo, Japan) for 13 minutes.

### Glycosylation staining

Glycosylation of Pfs230 domains was assessed using the Pierce Glycoprotein Staining Kit (ThermoFisher Scientific). 5 μg of each Pfs230 protein construct was loaded on gel and the gel was stained following manufacturer’s instructions.

### Mass spectrometry

The identity of recombinant Pfs230D12 was confirmed by mass spectrometry. 5 μg of purified protein was denatured in 4M urea, 100 mM Tris-HCl (pH 8.0) and disulfide bonds were reduced using 10 mM DTT for 30 minutes at RT. Cysteines were alkylated using 50 mM Iodoacetamide for 30 minutes, and samples were diluted to 2M Urea, using 100 mM Tris-HCl (pH 8.0). Samples were digested overnight with 0.5 μg Trypsin at 25°C. Next day, samples were desalted using StageTips ^31^.

Peptides were analyzed using an Easy nLC 1000 equiped with a 30 cm reverse phase column, coupled on-line to an Orbitrap Fusion Tribrid mass spectrometer (Thermo Scientific). A 60 minute gradient of buffer B (80% acetonitrile, 0.1% formic acid) was applied and the mass spectrometer was operated in TopS mode with a dynamic exclusion of 60 seconds.

RAW data was analyzed using Maxquant ^32^ version 1.6.6.0 with a *Drosophila* database supplemented with sequences for single domain constructs of Pfs230. The mass spectrometry proteomics data have been deposited to the ProteomeXchange Consortium via the PRIDE partner repository with the dataset identifier PXD039716 ^33^.

### Mice immunization

45 female 6-8 weeks old CD-1 mice (Charles Rivers), divided in groups of 5 mice, were immunized with 230CMB, D1, D5, D6, D8, D9, D10, D12 and D13. 230CMB, a construct comprising aa 444-730 of Pfs230 produced in a plant-based expression system ^21^ and known to induce transmission reducing capacity was included as positive control. Mice were injected subcutaneously with 100 μL of 0.2 mg/mL antigen in 70% Montanide ISA720 (SEPPIC) at day 0, day 21 and day 43. Pre-bleed samples were collected at day -1 and final bleeding was performed at day 57 before sacrificing (Figure 2A). Blood was allowed to clot at RT for 30 minutes, and serum was collected after centrifugation and was stored at -20°C. For each group of mice sera, samples were pooled for further analysis.

### Enzyme-Linked Immunosorbent Assays (ELISA)

For antigen ELISA to assess antigen-specific antibody responses in mice, Nunc MaxoSorp 96-wells plates (ThermoFisher) were coated with 100 μL of 1 μg/mL antigen and incubated overnight at 4°C. Plates were washed three times with PBS in between incubation steps. Plates were blocked with 5% mPBS for 1 hour. Plates were incubated with serum samples diluted in 1% mPBST, for 3 hours at RT. Subsequently, the plates were incubated with polyclonal Rabbit Anti-Mouse HRP (1/3000 dilution, Cat. No. P0260, DAKO) for 2 hours at RT. The ELISA was developed by adding 100 μL tetramethylbenzidine (TMB). The color reaction was stopped by adding 50 μL 0.2 M H_2_SO_4_ and the optical density was read at 450 nm on an iMark™ microplate absorbance reader (Bio-Rad).

For the gametocyte ELISA, *P. falciparum* NF54 gametocyte extracts were prepared as described previously ^30^. 100 μL lysate per well, equivalent to 75,000 gametocytes, was pipetted into Nunc MaxiSorp™ 96-wells plates (ThermoFisher) and incubated overnight at 4°C. The next steps of the ELISA were performed as described above.

D12-specific antibody levels in sera from individuals exposed to malaria parasites were assessed using an antigen ELISA as described above. Goat anti-Human IgG (H+L) HRP (1/40,000 dilution, Cat. No. 31412, Invitrogen) was used for detection. A titration of pooled hyperimmune serum from gametocyte carriers in Tanzania was used to calculate arbitrary units using ADAMSEL FPL (http://www.malariaresearch.eu/content/sopware).

### Gamete purification

To obtain purified female gametes for flow cytometry assays, we collected N-acetyl glucosamine treated 16-day old *P. falciparum* NF54 gametocyte cultures. The cultures were centrifuged for 10 minutes at 2,000×g at RT, to be resuspended in FBS using a volume that equals half the original culture volume. Gametocytes were placed on a roller bank for 45 minutes at RT for activation and thereafter centrifuged for 10 minutes at 2,000×g at 4°C. The pelleted gametes were resuspended in 1 mL PBS, loaded onto a 7 mL layer of 11% w/v Accudenz (Accurate Chemical) and centrifuged for 30 minutes at 7,000×g at 4°C without brake (Sorvall RC-5B Superspeed Centrifuge with HB-4 swing-out rotor). The female gametes present in the top layer were collected, transferred to a 50 mL tube and PBS was added up to 50 mL total volume. A final centrifugation for 5 minutes at 2,000×g at 4°C was done to pellet the gametes, which were resuspended in 1 mL PBS and counted using a Bürker-Turk counting chamber.

### Flow cytometry

For the assessment of gamete antibody binding, gamete C1q deposition and gamete lysis we used similar flow cytometry assays with specific adjustments that are described in this paragraph. All gamete incubations were carried out in PBS supplemented with 2% FBS and 0.02% sodium azide. For all three assays 50,000 purified gametes were used per well in a V-bottom non-treated 96-well plate (Costar) and were incubated for 1 hour at RT with mice sera. In the case of a lysis assay, there is an addition of 20% normal human serum (NHS) and the incubation is reduced to 30 minutes at RT. For a C1q deposition assay this is reduced to 10% NHS (30 minutes incubation at RT). Plates were centrifuged at 2,000×g for 3 minutes at 4°C and washed three times with PBS. The C1q deposition assay includes additional steps; first PBS supplemented with 10 mM EDTA is added for 5 minutes at 4°C to inactivate complement. Second, after 3 washes with PBS, 1/5000 anti-C1q goat anti-human polyclonal serum (Complement Technology) is added for 30 minutes incubation at RT. Gametes were washed and then incubated with either 1/200 Alexa Fluor™ 488 Chicken anti-Mouse IgG (H+L) (Invitrogen) (binding assay), 1/200 anti-Pfs47 (rat mAb 47.1) ^34^ labelled with DyLight™ 650 NHS ester (Thermo Scientific) (lysis assay) or 1/200 Alexa Fluor™ 488 Donkey anti-Goat IgG (H+L) (Invitrogen). 1/1000 eBioscience™ Fixable Viability Dye eFluor™ 780 (Invitrogen) was added in all assays and gametes were incubated for 30 minutes at RT. After washing with PBS, samples were resuspended in 150 μl PBS. Antibody binding to gametes, lysis of gametes and C1q deposition on gametes were assessed by flow cytometry by analysing a minimum of 2,000 gametes with the Gallios™ 10-color system (Beckman Coulter) and analyzed with FlowJo (BD, version 10.7.1) (gating strategy in Supplementary figure 4).

### SMFA

Mice sera were diluted in FCS and mixed with mature *P. falciparum* NF54 gametocytes and NHS. To inactivate NHS for conditions where inactive complement is required, it was heated for 30 minutes at 56°C prior to mixing with the gametocytes. *Anopheles stephensi* mosquitoes from a colony maintained at Radboudumc (Nijmegen, the Netherlands) were fed blood meals as described previously ^35^. Unfed and partially fed mosquitoes were removed. 20 mosquitoes per condition were dissected 6-8 days after the blood meal to collect their midguts. The midguts were stained with mercurochrome and oocysts were counted. TRA was defined as the reduction in oocyst intensity (oocysts per mosquito midgut) in a test condition compared to a negative control in which no mice sera (FBS control) was added. All samples were tested in two independent SMFA experiments for which the data are shown in Supplementary file 1.

### DMFA

Gametocyte-infected blood from patients residing in the villages surrounding Bobo-Dioulasso was collected in heparin tubes, 5 mL per tube. Immediately after blood collection, the blood was centrifuged at 3000×g for 5 minutes, and plasma was removed. 120 μl of the remaining RBC pellet was transferred to tubes containing 90 μl of naïve AB serum and 30 μl of mice sera (or FCS as negative control). The total of 240 μl was carefully mixed by pipetting and the content of each tube was transferred to an individual feeder maintained at 37°C to allow *Anopheles coluzzii* mosquito feeding for 30 minutes. Unfed mosquitoes were removed; fullyfed mosquitoes were kept for 7 days post feeding. All surviving mosquitoes were dissected for each condition, midguts were stained with 0.5% mercurochrome for oocyst detection and oocysts were counted.

### Statistical analysis

Transmission reducing activity (TRA) was calculated as the reduction in oocysts compared to a negative control, using an online tool ^36^. All other statistical analyses were performed using GraphPad Prism (version 10.1.0).

## Supporting information

Supplementary Tables Raw SMFA and DMFA Data

## Ethical statement

Asymptomatic gametocyte carriers, aged 7 and 10 years, were enrolled in November 2023 from the villages surrounding Bobo Dioulasso (Burkina Faso). Venous blood samples of the two volunteers were collected after written informed consent was obtained from participants or their guardian(s). Ethical approval was provided by the Ethical Review Committee of the Ministry of Health, Burkina Faso (N°2022-05-093); Institutional ethics review committee for health science research Bobo Dioulasso (A014-2022-CEIRES). For the cohort study in Uganda, ethical approval was obtained from the Makerere University School of Medicine Research and Ethics Committee, the Uganda National Council for Science and Technology, the London School of Hygiene & Tropical Medicine Ethics Committee, the Durham University School of Biological and Biomedical Sciences Ethics Committee, and the University of California, San Francisco, Committee on Human Research.

### Acknowledgements

We would like to thank the Laura Pelser-Posthumus, Astrid Pouwelsen, Jacqueline Kuhnen, Jolanda Klaassen and Saskia Mulder for mosquito dissections, and Team 2 of the Animal Research Facility (CDL) at Radboudumc for performing mouse immunizations. We are grateful to the study participants who participated in the longitudinal study in Uganda. We are grateful to the volunteers from Bobo Dioulasso for participating in this study and our colleagues at IRSS for conducting DMFA experiments. We would like to thank Sara Lynn Blanken and Will Stone for assistance with analysis of naturally acquired immune responses to Pfs230D12, and Jessica Chichester (Fraunhofer) for kindly providing us the 230CMB protein construct.

## Supplementary files

Supplementary file 1 (SMFA and DMFA data).

