## Supplementary Figures and Tables for "Pfs230 Domain 12 is a potent malaria transmission-blocking vaccine candidate"

### Supplemental Information
### Tables and figures

Maartje R. Inklaar[1], Roos M. de Jong[1], Dari F. Da[2], Lisanne L. Hubregtse[1], Maartje Meijer[1], Karina Teelen[1], Marga van de Vegte-Bolmer[1], Geert-Jan van Gemert[1], Rianne Stoter[1], Hikaru Nagaoka[3], Takafumi Tsuboi[4], Eizo Takashima[3], Cornelia G. Spruijt[5], Michiel Vermeulen[5], Roch K. Dabire[2], Emmanuel Arinaitwe[6], Anna Cohuet[7], Teun Bousema[1], Matthijs M. Jore[1,*]

[1] *Department of Medical Microbiology, Radboud University Medical Center, Nijmegen, The Netherlands*
[2] *Institut de Recherche en Sciences de la Santé, Direction Régionale, Bobo Dioulasso, Burkina Faso*
[3] *Division of Malaria Research, Proteo-Science Center, Ehime University, Matsuyama, Japan*
[4] *Division of Cell-Free Sciences, Proteo-Science Center, Ehime University, Matsuyama, Japan*
[5] *Department of Molecular Biology, Faculty of Science, Oncode Institute, Radboud University Nijmegen, Nijmegen, The Netherlands*
[6] *Infectious Diseases Research Collaboration, Kampala, Uganda*
[7] *MIVEGEC, Montpellier University, IRD, CNRS, Montpellier, France*

**Supplementary table 1. Overview of Pfs230 single domain sequences as they are expressed in S2 cells.**

| **Pfs230 single domain** | **Nucleotide sequence**<br>Underlined = kozak sequence<br>*Italic* = signal peptide<br>**Blue = His-tag**<br>**Dark grey = linker (also restriction site Ngo MIV)**<br>**Grey = domain**<br>**Yellow = linker**<br>**Dark blue = C-tag** |
|---|---|
| D1 | GCCACC*ATGAAGCTGTGCATCCTGCTGGCCGTGGTGGCCTTCGTGGGACTGAGCCTGGGA*CACC<br>ACCACCATCACCACGCCGGCAAAGAGTACGTCTGCGACTTCACCGATCAGCTGAAGCCAACCGA<br>GTCGGGCCCCAAAGTGAAGAAATGCGAAGTGAAAGTGAACGAGCCCCTGATCAAAGTCAAGATT<br>ATCTGCCCGCTGAAGGGCAGCGTGGAAAAGCTGTACGATAACATCGAGTACGTGCCCAAGAAAA<br>GCCCCTACGTGGTGCTGACCAAAGAGGAAACGAAGCTGAAAGAGAAGCTGCTGAGCAAGCTGAT<br>CTACGGCCTGCTGATCTCCCCGACCGTGAACGAGAAAGAGAACAACTTCAAAGAGGGCGTCATC<br>GAGTTCACCCTGCCGCCAGTGGTGCATAAGGCCACCGTGTTCTACTTCATCTGCGACAACAGCA<br>AGACCGAGGACGATAACAAGAAGGGCAACCGCGGCATCGTGGAAGTGTACGTGGAACCCTACGG<br>ATCAGAGCCCGAGGCCTAA |
| D2 | GCCACC*ATGAAGCTGTGCATCCTGCTGGCCGTGGTGGCCTTCGTGGGACTGAGCCTGGGA*CACC<br>ACCACCATCACCACGCCGGCGGCAACAAGATCAACGGCTGCGCCTTCCTGGATGAGGATGAGGA<br>AGAAGAGAAGTACGGCAATCAGATCGAAGAGGACGAGCACAACGAGAAGATCAAGATGAAGACC<br>TTCTTCACCCAAAACATCTACAAGAAGAACAACATCTACCCGTGCTACATGAAGCTGTACTCCG<br>GCGATATCGGCGGCATTCTGTTCCCCAAGAACATCAAGAGCACGACCTGCTTCGAGGAAATGAT<br>CCCCTACAACAAAGAAATCAAGTGGAACAAAGAGAACAAGAGCCTGGGCAACCTGGTCAACAAC<br>AGCGTGGTGTATAACAAAGAGATGAACGCCAAGTACTTCAACGTGCAGTACGTGCACATCCCCA<br>CCAGCTACAAGGATACCCTGAACCTGTTCTGCAGCATCATCCTGAAAGAGGAAGAGAGCAACCT<br>GATCAGCACCTCCTACCTGGTGTACGTGTCCATCAACGAGGGATCAGAGCCCGAGGCCTAA |
| D3 | GCCACC*ATGAAGCTGTGCATCCTGCTGGCCGTGGTGGCCTTCGTGGGACTGAGCCTGGGA*CACC<br>ACCACCATCACCACGCCGGCCACGATTATACCTGCGATTTCACGGACAAGCTCGACAAGACCGT<br>GCCGAGCACCGCCAATGGCAAGAAGCTGTTCATCTGCCGCAAGCACCTGAAAGAATTCGACACC<br>TTCACGCTGAAGTGCAACGTGAACAAGACGCAGTACCCCAACATCGAGATCTTCCCAAAGACGC<br>TGAAGGACAAGAAAGAGGTCCTGAAGCTGGATCTGGACATCCAGTACCAGATGTTCAGCAAGTT<br>CTTCAAGTTTAACACCCAGAACGCGAAGTACCTGAATCTGTACCCCTACTACCTGATCTTCCCC<br>TTCAACCACATCGGAAAGAAAGAGCTGAAAAACAACCCCACCTACAAGAACCACAAGGACGTGA<br>AGTATTTCGAGCAGTCCTCCGTGCTGAGCCCACTGAGTAGTGCCGATAGCCTGGGAAAGCTGTT<br>GAACTTCCTGGACACCCAAGAGACAGTGTGCCTGACCGAGAAGATTCGCTATCTGAACCTGAGC<br>ATCAATGAGCTGGGCAGCGATAACAACACCTTCTCCGTGACGTTCCAGGTGCCGCCGTACATCG<br>ATATCAAAGAACCCTTCTACTTTATGTTCGGCTGCAACAACAACAAAGGCGAGGGCAACATCGG<br>CATAGTCGAGCTGCTGATTAGCAAGCAAGGATCAGAGCCCGAGGCCTAA |
| D4 | GCCACC*ATGAAGCTGTGCATCCTGCTGGCCGTGGTGGCCTTCGTGGGACTGAGCCTGGGA*CACC<br>ACCACCATCACCACGCCGGCGAAGAAAAGATTAAGGGCTGCAATTTCCACGAGTCCAAGCTGGA<br>CTACTTCAATGAGAACATCAGCAGCGATACCCACGAGTGCACGCTGCACGCCTATGAGAACGAT<br>ATCATCGGCTTCAACTGCCTGGAAACGACGCACCCCAACGAGGTGGAAGTGGAAGTTGAGGATG<br>CCGAGATCTATCTGCAGCCCGAGAACTGCTTCAACAACGTCTACAAGGGCCTGAACTCCGTGGA<br>TATCACCACCATCCTGAAGAACGCCCAGACCTACAACATTAACAACAAAAAGACCCCGACCTTC<br>CTGAAGATCCCGCCATACAACCTGCTGGAAGATGTGGAAATCAGCTGCCAGTGCACCATCAAGC<br>AAGTGGTCAAGAAAATCAAAGTGATCATCACCAAGAACGACGGATCAGAGCCCGAGGCCTAA |
| D5 | GCCACC*ATGAAGCTGTGCATCCTGCTGGCCGTGGTGGCCTTCGTGGGACTGAGCCTGGGA*CACC<br>ACCACCATCACCACGCCGGCAAGATCTACAAGTGCGAGCACGAGAACTTCATCAACCCGCGCGT<br>CAACAAGACCTTCGACGAGAACGTCGAGTACACGTGCAATATCAAGATCGAGAATTTCTTCAAC<br>TACATCCAGATTTTCTGCCCCGCCAAGGATCTGGGCATCTATAAGAATATCCAGATGTACTACG<br>ACATCGTGAAGCCGACGCGCGTGCCCCAGTTCAAAAAATTCAACAATGAGGAGCTCCACAAGCT<br>CATCCCCAACTCCGAGATGCTGCACAAGACGAAAGAGATGCTGATCCTGTACAACGAAGAGAAG<br>GTGGACCTGCTGCACTTCTACGTGTTCTTGCCCATCTACATCAAGGACATCTACGAGTTCAACA<br>TCGTGTGCGACAACTCCAAGACGATGTGGAAGAACCAGCTCGGCGGAAAAGTGATCTACCACAT |

| | |
|---|---|
| | CACCGTCAGCAAGCGCGGATCAGAGCCCGAGGCCTAA |
| D6 | GCCACC*ATGAAGCTGTGCATCCTGCTGGCCGTGGTGGCCTTCGTGGGACTGAGCCTGGGA*CACC<br>ACCACCATCACCACGCCGGCTTCGATAACGAGCACGCCCACATGTTCAGCTATAACAAGACCAA<br>CGTGAAGAACTGCATCATCGACGCCAAGCCGAAGGATCTGATCGGCTTCGTGTGCCCAAGCGGC<br>ACACTGAAGCTGACCAATTGCTTCAAGGATGCCATCGTGCACACCAACCTGACCAACATCAACG<br>GCATCCTGTATCTCAAGAACAACCTGGCCAACTTCACGTACAAGCACCAGTTCAATTACATGGA<br>AATCCCCGCGCTGATGGACAACGACATCAGCTTCAAGTGCATCTGCGTGGACCTGAAGAAAAAG<br>AAGTACAACGTCAAGAGCCCGCTGGGCCCCGGATCAGAGCCCGAGGCCTAA |
| D7 | GCCACC*ATGAAGCTGTGCATCCTGCTGGCCGTGGTGGCCTTCGTGGGACTGAGCCTGGGA*CACC<br>ACCACCATCACCACGCCGGCAACCGCCACGTGTGCGATTTCTCCAAGAACAATCTGATCGTGCC<br>CGAGTCGTTGAAGAAGAAAGAGGAACTCGGCGGCAACCCCGTGAACATCCATTGCTATGCCCTG<br>TTGAAGCCCCTGGATACGCTGTATGTGAAGTGCCCCACCTCCAAGGATAACTACGAGGCCGCCA<br>AAGTCAACATCAGCGAGAATGATAACGAGTACGAGTTGCAAGTGATCTCCCTGATCGAGAAGCG<br>CTTTCACAACTTCGAGACACTGGAAAGCAAAAAGCCCGGCAACGGCGACGTCGTGGTGCACAAT<br>GGTGTTGTGGATACCGGACCGGTGCTGGATAACTCCACGTTCGAGAAGTACTTTAAGAACATTA<br>AGATCAAGCCCGATAAGTTCTTCGAGAAAGTTATCAATGAGTACGACGACACCGAGGAAGAAAA<br>GGACCTGGAATCCATCCTGCCAGGCGCCATCGTGTCCCCAATGAAGGTGCTCAAGAAGAAGGAC<br>CCCTTCACCAGCTATGCCGCCTTTGTGGTGCCACCGATCGTGCCAAAGGATCTGCACTTCAAGG<br>TGGAATGCAACAATACCGAGTACAAGGACGAGAACCAGTACATCAGCGGCTACAATGGCATCAT<br>CCACATCGACATCTCCAACAGCGGATCAGAGCCCGAGGCCTAA |
| D8 | GCCACC*ATGAAGCTGTGCATCCTGCTGGCCGTGGTGGCCTTCGTGGGACTGAGCCTGGGA*CACC<br>ACCACCATCACCACGCCGGCAACCGCAAGATCAATGGATGCGACTTTAGCACCAACAACTCCAG<br>CATCCTGACCAGCTCCGTGAAGCTGGTTAACGGCGAGACAAAGAACTGCGAGATCAATATCAAC<br>AACAACGAAGTGTTCGGCATCATCTGTGACAATGAGACAAATCTGGACCCAGAGAAGTGCTTCC<br>ATGAGATCTACTCCAAGGACAACAAGACGGTCAAAAAGTTCCGCGAAGTGATCCCCAATATCGA<br>CATTTTCAGCCTGCACAACTCGAACAAGAAAAAGGTGGCCTACGCCAAGGTGCCCCTGGACTAT<br>ATTAACAAGCTGCTGTTCAGCTGCTCCTGCAAGACCAGCCACACCAACACCATCGGCACGATGA<br>AAGTGACCCTGAACAAAGACGAGGGATCAGAGCCCGAGGCCTAA |
| D9 | GCCACC*ATGAAGCTGTGCATCCTGCTGGCCGTGGTGGCCTTCGTGGGACTGAGCCTGGGA*CACC<br>ACCACCATCACCACGCCGGCAACGTGCACCTGTGCAATTTCTTCGACAACCCCGAGCTGACCTT<br>CGACAACAACAAGATCGTGCTGTGCAAGATCGATGCCGAGCTGTTTAGCGAAGTCATCATTCAG<br>CTGCCCATCTTCGGCACCAAAAACGTCGAGGAAGGCGTCCAGAACGAAGAGTACAAGAAGTTCA<br>GCCTGAAGCCGAGCCTGGTGTTCGATGATAACAACAATGACATCAAAGTCATCGGCAAAGAGAA<br>GAACGAGGTTTCCATCTCGCTGGCCCTGAAGGGCGTGTACGGCAACCGCATCTTCACCTTTGAT<br>AAGAACGGCAAGAAAGGCGAAGGCATCAGCTTTTTCATCCCGCCGATCAAGCAGGATACCGACC<br>TGAAGTTTATCATCAACGAAACCATCGATAACAGCAACATTAAGCAGCGCGGCCTGATCTACAT<br>CTTCGTGCGCAAGAACGTGTCGGGATCAGAGCCCGAGGCCTAA |
| D10 | GCCACC*ATGAAGCTGTGCATCCTGCTGGCCGTGGTGGCCTTCGTGGGACTGAGCCTGGGA*CACC<br>ACCACCATCACCACGCCGGCGAGAACTCGTTCAAGCTGTGTGATTTCACCACCGGCAGCACCAG<br>CCTGATGGAATTGAACAGCCAAGTGAAAGAAAAGAAGTGCACCGTTAAGATTAAGAAGGGCGAT<br>ATCTTCGGCCTGAAATGCCCCAAGGGATTCGCCATTTTTCCGCAAGCCTGCTTCTCCAACGTCC<br>TGCTCGAGTACTACAAGAGCGATTACGAGGACAGCGAGCACATCAACTACTACATTCACAAGGA<br>CAAAAAGTACAATCTGAAGCCCAAGGACGTTATCGAGTTGATGGATGAGAACTTCCGCGAGCTG<br>CAAAACATTCAGCAGTACACCGGCATCAGCAACATCACCGATGTGCTGCATTTCAAGAACTTCA<br>ACCTGGGCAATCTGCCGCTCAACTTCAAGAATCACTACAGCACCGCCTATGCGAAGGTGCCGGA<br>TACCTTCAACTCCATCATCAACTTCAGCTGCAACTGCTACAATCCCGAGAAGCACGTCTACGGC<br>ACCATGCAGGTCGAGAGCGATAACGGATCAGAGCCCGAGGCCTAA |
| D11 | GCCACC*ATGAAGCTGTGCATCCTGCTGGCCGTGGTGGCCTTCGTGGGACTGAGCCTGGGA*CACC<br>ACCACCATCACCACGCCGGCAATGAGCACATTTGCGACTACGAAAAGAACGAGTCGCTGATCTC<br>GACCCTGCCAAACGACACCAAGAAGATCCAGAAGTCGATCTGCAAGATTAACGCGAAGGCCCTG<br>GATGTGGTCACCATTAAGTGCCCGCATACCAAGAATTTCACCCCGAAGGATTACTTCCCCAACA<br>GCAGCCTGATCACCAACGATAAGAAGATCGTCATCACGTTCGATAAGAAAAACTTCGTCACCTA<br>CATCGACCCCACCAAAAAGACGTTCTCCCTGAAAGACATCTACATTCAGAGCTTCTACGGCGTG<br>TCCCTGGATCACCTGAACCAGATCAAAAAAATCCACGAGGAATGGGACGACGTCCACCTGTTTT |

| | |
|---|---|
| | ACCCGCCGCACAACGTTCTGCACAACGTGGTCCTGAACAACCACATTGTGAACCTGTCCAGCGC<br>CTTGGAGGGCGTGCTGTTCATGAAGTCCAAAGTGACCGGCGACGAGACAGCCACGAAGAAGAAT<br>ACCACACTGCCCACCGATGGCGTGTCCAGCATTCTGATCCCGCCGTACGTGAAAGAAGATATCA<br>CCTTCCATCTGTTCTGCGGCAAGTCCACGACCAAGAAGCCCAACAAAAAAAACACCAGCTTGGC<br>CCTGATCCACATTCACATCAGCTCCAATGGATCAGAGCCCGAGGCCTAA |
| D12 | GCCACC*ATGAAGCTGTGCATCCTGCTGGCCGTGGTGGCCTTCGTGGGACTGAGCCTGGGA*CACC<br>ACCACCATCACCACGCCGGCCGCAATATCATCCACGGCTGCGACTTTCTGTACCTGGAAAACCA<br>GACCAACGACGCCATCTCGAACAACAACAACAACTCCTACAGCATCTTCACCCACAACAAGAAC<br>ACCGAGAACAACCTCATCTGCGATATTTCGCTGATCCCCAAGACCGTGATCGGCATCAAGTGCC<br>CCAACAAGAAGCTGAACCCGCAGACCTGCTTTGACGAGGTGTACTACGTCAAACAAGAGGACGT<br>GCCGTCCAAGACCATCACCGCCGACAAGTACAATACCTTCAGCAAGGATAAGATTGGCAACATC<br>CTCAAAAACGCCATCAGCATCAACAACCCGGACGAGAAGGATAACACCTACACCTATCTGATCC<br>TGCCGGAAAAGTTCGAGGAAGAGTTGATCGATACCAAAAAGGTGCTGGCCTGCACGTGTGACAA<br>CAAGTACATTATCCACATGAAGATCGAAAAGTCCACCGGATCAGAGCCCGAGGCCTAA |
| D13 | GCCACC*ATGAAGCTGTGCATCCTGCTGGCCGTGGTGGCCTTCGTGGGACTGAGCCTGGGA*CACC<br>ACCACCATCACCACGCCGGCGGCAAGGATATCTGCAAATACGACGTGACCACCAAGGTGGCCAC<br>GTGCGAGATTATCGACACCATCGATTCGAGCGTGCTGAAAGAACACCACACCGTGCACTACTCG<br>ATCACCCTGTCGCGCTGGGATAAGCTGATCATCAAGTACCCGACCAACGAGAAAACCCACTTTG<br>AGAACTTTTTCGTGAACCCGTTCAACCTCAAGGACAAGGTGCTCTACAATTACAACAAGCCCAT<br>CAACATTGAGCACATACTGCCCGGTGCCATCACCACCGATATCTACGATACGCGCACCAAGATT<br>AAGCAGTACATCCTGCGCATCCCACCGTATGTGCACAAGGATATTCACTTCTCCCTGGAATTCA<br>ACAACTCCCTGAGCCTGACCAAGCAGAACCAGAACATTATCTACGGCAATGTGGCCAAGATCTT<br>CATCCATATCAACCAGGGCGGATCAGAGCCCGAGGCCTAA |
| D14 | GCCACC*ATGAAGCTGTGCATCCTGCTGGCCGTGGTGGCCTTCGTGGGACTGAGCCTGGGA*CACC<br>ACCACCATCACCACGCCGGCTACAAAGAGATCCACGGTTGCGATTTCACCGGCAAGTACAGCCA<br>CCTGTTCACCTACTCCAAAAAGCCGCTGCCGAACGATGACGACATCTGCAATGTGACCATCGGA<br>AACAACACGTTCAGCGGATTCGCCTGCCTGTCGCACTTCGAGCTGAAACCCAACAACTGCTTCT<br>CGTCGGTGTACGATTACAACGAGGCCAACAAAGTGAAAAAGTTGTTCGACCTGTCGACCAAGGT<br>CGAGCTGGATCACATCAAACAGAACACCTCCGGCTACACCCTGTCCTACATCATTTTTAACAAA<br>GAATCGACCAAGCTCAAGTTCTCCTGCACATGCAGCAGCAACTACTCCAACTACACCATCCGCA<br>TCACCTTCGATCCGGGATCAGAGCCCGAGGCCTAA |

**Supplementary table 2. Characteristics of all Pfs230 single domain constructs.** NetOGlyc version 4.0 (DTU Health Tech, Denmark) [1] and NetNglyc version 1.0 (DTU Health Tech, Denmark) [2] were used to predict the O-linked and N-linked glycosylation sites.

| Pfs230 single Domain | Amino acid numbers within full-length Pfs230 | Molecular weight (kDa) | Number of cysteines | Predicted N-linked glycosylation sites | Predicted O-linked glycosylation sites |
|---|---|---|---|---|---|
| D1 | 589-730 | 17.8 | 4 | 0 | 2 |
| D2 | 731-886 | 19.8 | 4 | 2 | 0 |
| D3 | 918-1133 | 26.8 | 5 | 4 | 2 |
| D4 | 1134-1268 | 17.1 | 6 | 1 | 0 |
| D5 | 1285-1432 | 19.5 | 4 | 1 | 0 |
| D6 | 1433-1550 | 14.1 | 6 | 2 | 0 |
| D7 | 1694-1907 | 25.9 | 4 | 3 | 1 |
| D8 | 1908-2036 | 16.1 | 6 | 4 | 1 |
| D9 | 2052-2201 | 18.7 | 2 | 1 | 0 |
| D10 | 2202-2373 | 21.5 | 6 | 2 | 0 |
| D11 | 2448-2663 | 26.1 | 4 | 7 | 4 |
| D12 | 2664-2818 | 19.4 | 6 | 1 | 0 |
| D13 | 2831-2979 | 19.0 | 2 | 0 | 0 |
| D14 | 2980-3105 | 16.0 | 6 | 5 | 0 |

**Supplementary table 3. Mass spectrometry results confirm Pfs230-D12 identity**. The table includes all Pfs230 recombinant domains produced in this study that were found in the peptide preparation.

| Protein ID | Intensity | % of total intensity | Peptides | Razor + unique peptides | Unique peptides | % coverage |
|---|---|---|---|---|---|---|
| Pfs230-D12 | $9.853 \times 10^{11}$ | 98.226 | 29 | 29 | 29 | 89.9 |
| Pfs230-D5 | $5.715 \times 10^{6}$ | 0.001 | 2 | 2 | 2 | 15.6 |

**Supplementary table 4. Peptides, covering recombinant Pfs230 fragments, identified by mass spectrometry.**

| Domain | Peptide |
|---|---|
| D12 | CPNKKLNPQTCFDEVYYVK |
| D12 | DKIGNILK |
| D12 | DNTYTYLILPEK |
| D12 | DNTYTYLILPEKFEEELIDTK |
| D12 | DNTYTYLILPEKFEEELIDTKK |
| D12 | FEEELIDTK |
| D12 | FEEELIDTKK |
| D12 | KLNPQTCFDEVYYVK |
| D12 | KLNPQTCFDEVYYVKQEDVPSK |
| D12 | KVLACTCDNK |
| D12 | KVLACTCDNKYIIHMK |
| D12 | LNPQTCFDEVYYVK |
| D12 | LNPQTCFDEVYYVKQEDVPSK |
| D12 | NAISINNPDEK |
| D12 | NAISINNPDEKDNTYTYLILPEK |
| D12 | NAISINNPDEKDNTYTYLILPEKFEEELIDTK |
| D12 | NIIHGCDFLYLENQTNDAISNNNNNSYSIFTHNK |
| D12 | NTENNLICDISLIPK |
| D12 | NTENNLICDISLIPKTVIGIK |
| D12 | QEDVPSK |
| D12 | QEDVPSKTITADKYNTFSK |
| D12 | TITADKYNTFSK |
| D12 | TITADKYNTFSKDK |
| D12 | VLACTCDNK |
| D12 | VLACTCDNKYIIHMK |
| D12 | VLACTCDNKYIIHMKIEK |
| D12 | YIIHMKIEK |
| D12 | YNTFSKDK |
| D5 | IENFFNYIQIFCPAK |
| D5 | TFDENVEYTCNIK |

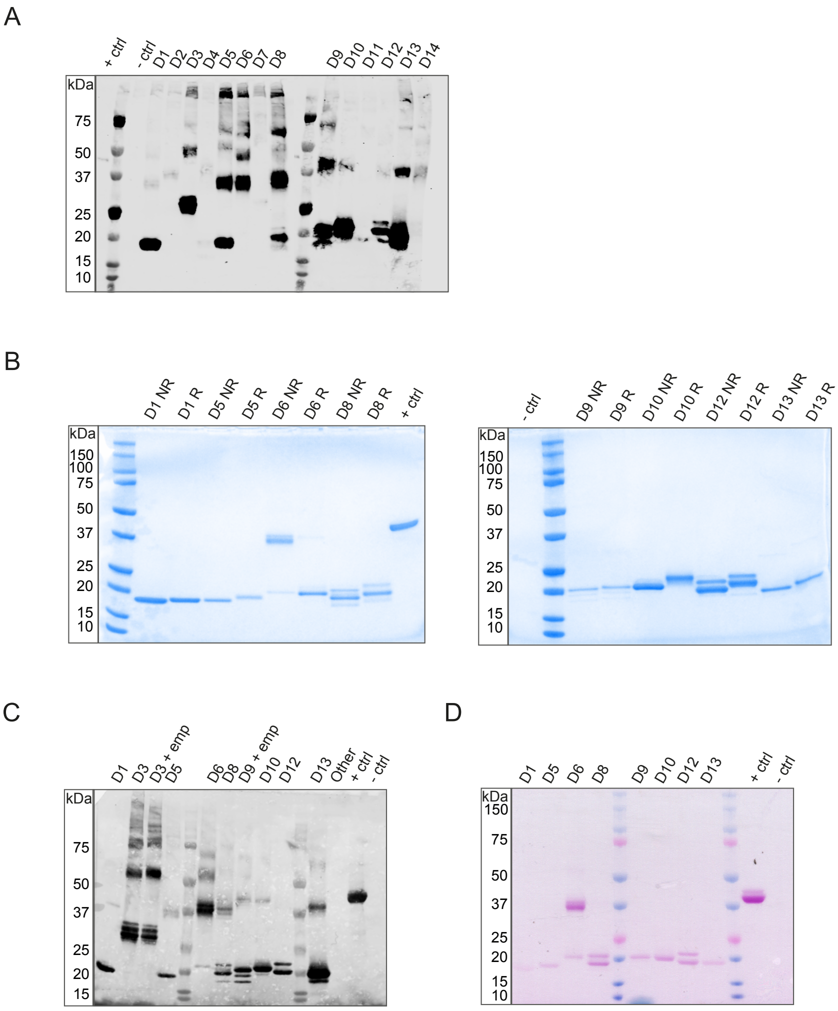


**Supplementary figure 1. Analysis of recombinant Pfs230 domain protein constructs produced in S2 cells. (A)** Western blot analysis of S2 cell supernatants. Supernatants of D1 till D14 were harvested from stable transfected cell lines. Supernatants were separated by SDS-PAGE, transferred to western blot, and stained with 1/1000 C-tag antibody and 1:2500 IRDye Streptavidin 680LT. + ctrl: Pro-CS3-6C; - ctrl: supernatant from un-transfected cells. **(B)** Coomassie-stained SDS-PAGE gels of Pfs230 single domain constructs purified from S2 cells and used for mice immunizations (original version of Figure 1B). NR: non-reduced conditions; R: reduced conditions. **(C)** Western blot of single domain Pfs230 constructs used for mice immunizations (original version of Figure 1C). **(D)** Glycoprotein staining after SDS-page (original version of Figure 1D).

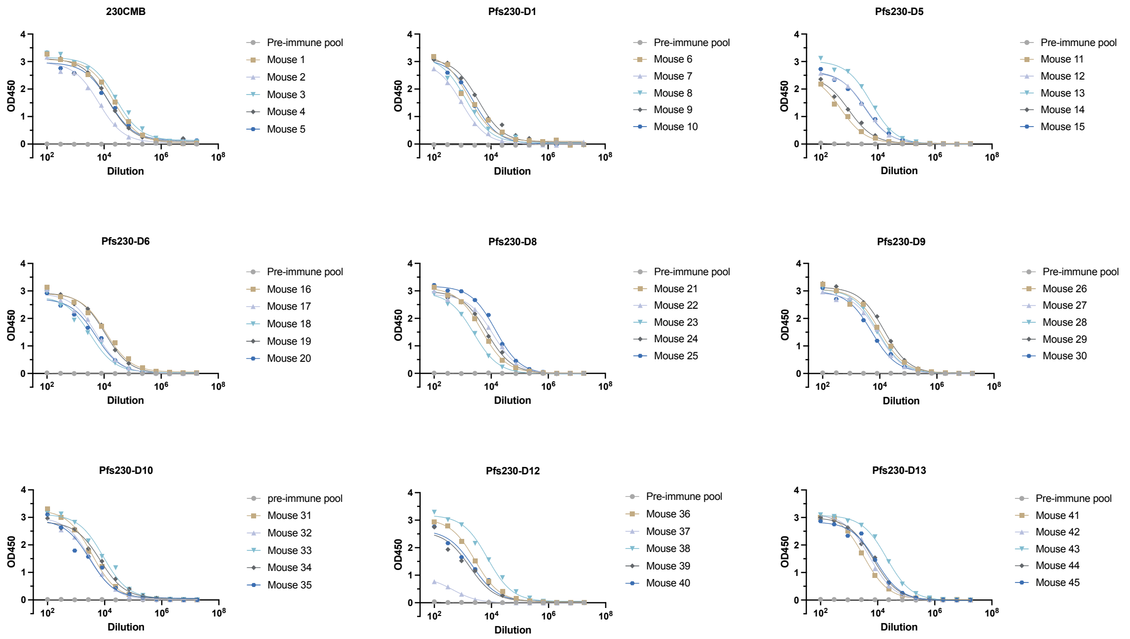


**Supplementary figure 2**. **Antigen specific ELISAs with sera from immunized mice.** The sera of each individual mouse were titrated (12-point titration, in singlicate) along with the pooled pre-immune sera. Sigmoidal curve fits were used to calculate $EC_{50}$ values that are shown in figure 1B.

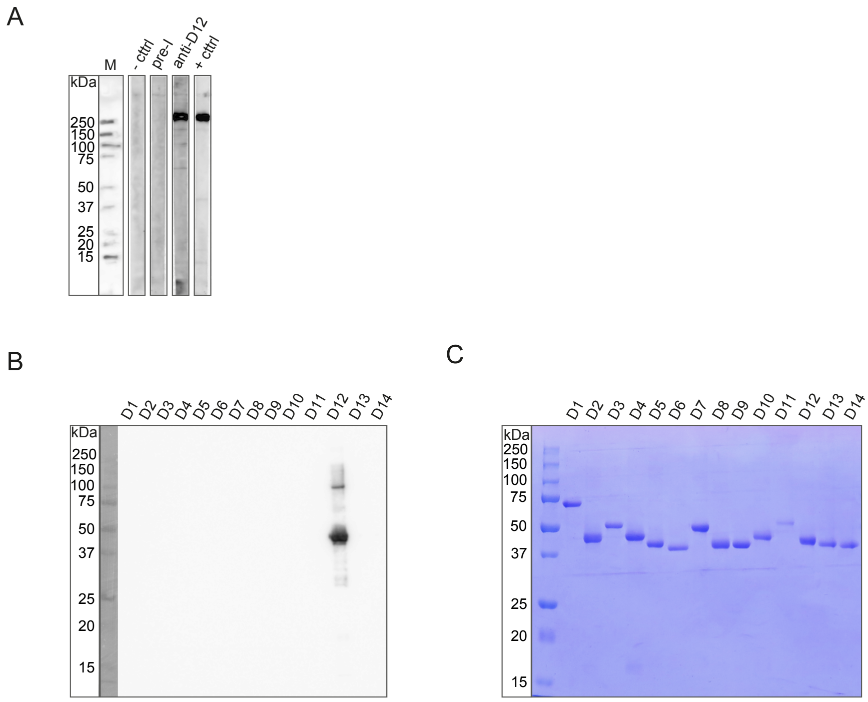


**Supplementary figure 3. Pfs230 and Pfs230-D12 specificity of the Pfs230-D12 induced antibodies.** Western blots incubated with mice sera induced by S2 cell-produced Pfs230-D12 showing specific recognition of **(A)** Pfs230 present in gametocyte extract and of **(B)** Pfs230-D12 recombinant protein produced by the wheat germ cell-free system. **(C)** The Pfs230 domains were separated by SDS-PAGE followed by Coomassie brilliant blue staining.

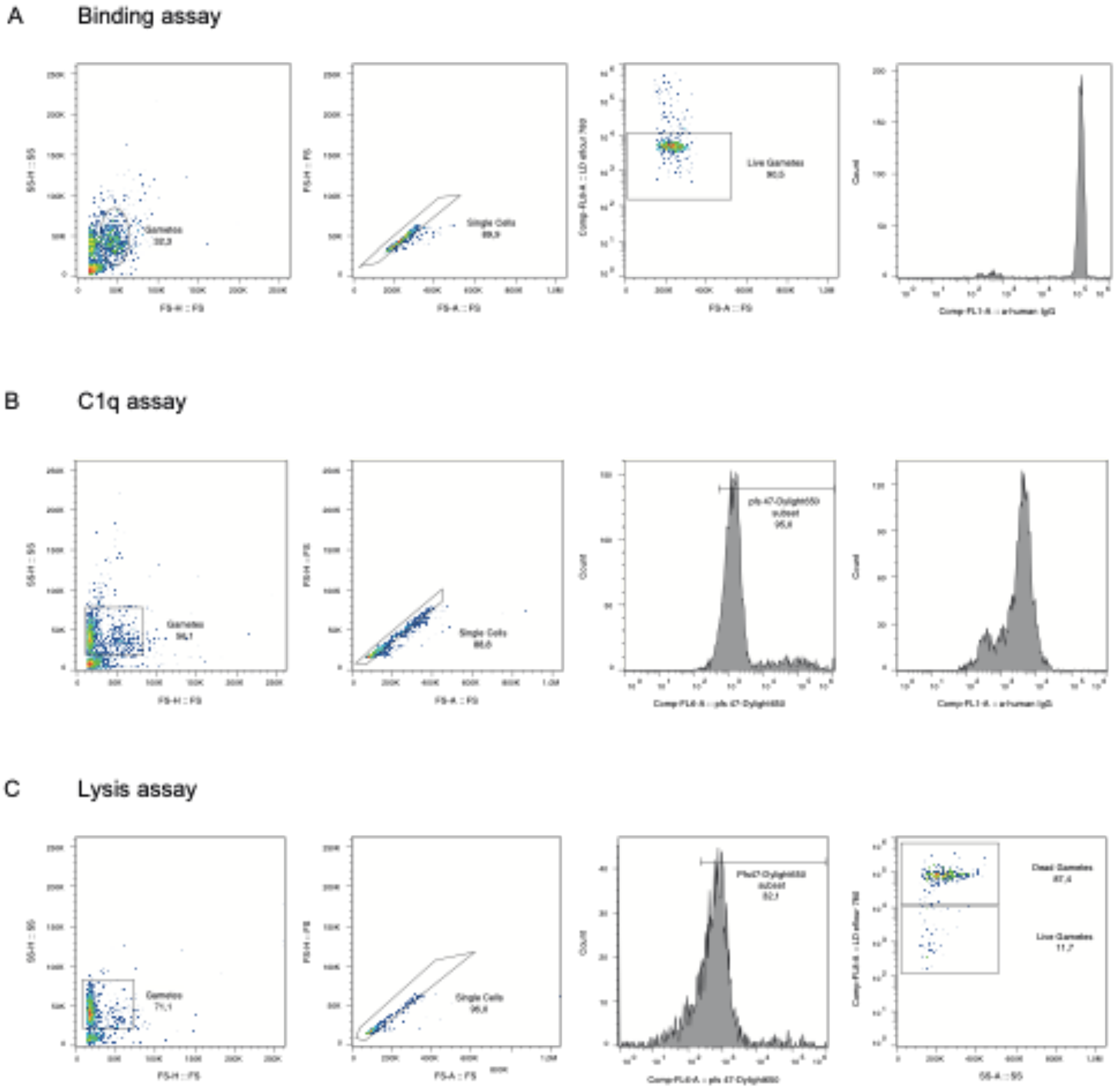


**Supplementary figure 4. Overview of flow cytometry gating strategies.** Exemplary plots that provide an overview of the gating strategy for **(A)** an antibody binding assay, **(B)** a C1q fixation assay and **(C)** a lysis assay, all with live female gametes. In the binding assay (A) gametes were gated for single cells (2nd column) and then live gametes were selected based on the absence of live-dead stain LD efluor 780, which stains dead cells (3rd column). For the C1q deposition assay **(B)** the gated single cells from the 2nd column were gated for gamete marker Pfs47 positivity, to then determine the anti-C1q deposition by FITC-labelled anti-C1q staining. For the lysis assay **(C)** gametes were gated for single cells (2nd column), and then for Pfs47 positivity. Dead gametes were stained with live-dead stain LD efluor 780 to determine the percentage dead cells (4th column). Note that two gamete populations can be observed in the forward scatter plots (1st column) in A and B; the left population contains dead gametes, the right population contains live gametes. In the antibody binding assay (A) and C1q deposition assay (B) we gated only the live population, while in the lysis assay (C) we gated both live and dead populations. SS-H = side scatter height, SS = side scatter, FS-H = forward scatter height, FS-A = forward scatter area, LD = live dead.
